# DisA limits RecA- and RadA/Sms-mediated replication fork remodelling to prevent genome instability

**DOI:** 10.1101/2020.11.23.394155

**Authors:** Rubén Torres, Juan C. Alonso

## Abstract

The DisA diadenylate cyclase (DAC), the DNA helicase RadA/Sms and the RecA recombinase are required to prevent a DNA replication stress during the revival of haploid *Bacillus subtilis* spores. Moreover, *disA, radA* and *recA* are epistatic among them in response to DNA damage. We show that DisA inhibits the ATPase activity of RadA/Sms C13A by competing for single-stranded (ss) DNA. In addition, DisA inhibits the helicase activity of RadA/Sms. RecA filamented onto ssDNA interacts with and recruits DisA and RadA/Sms onto branched DNA intermediates. In fact, RecA binds a reversed fork and facilitates RadA/Sms-mediated unwinding to restore a 3′-fork intermediate, but DisA inhibits it. Finally, RadA/Sms inhibits DisA DAC activity, but RecA counters this negative effect. We propose that RecA, DisA and RadA/Sms interactions, which are mutually exclusive, limit remodelling of stalled replication forks. DisA, in concert with RecA and/or RadA/Sms, indirectly contributes to template switching or lesion bypass, prevents fork breakage and facilitates the recovery of c-di-AMP levels to re-initiate cell proliferation.

**Subject Categories:** Genomic stability & Dynamics

## Introduction

Complete, accurate and timely DNA replication is essential to maintain genome integrity and cell proliferation. However, replicative DNA polymerases, which are generally poor at synthesising past lesions, are frequently hindered by obstacles, and replication stress occurs [1]. Cells have different mechanisms to prevent the incorrect handling of the perturbed replication forks and recombination functions to support replication fork movement [2-4]. Replication fork reversal, *i.e*., the active conversion of a stalled replication fork into a Holliday junction (HJ) structure, has emerged as a global and genetically controlled response to aid to the repair or bypass of DNA damage during replication stress [4-7].

In *Escherichia coli*, replication of DNA containing damaged template bases or DNA distortions can lead to spontaneous or remodeller-mediated fork reversal [4, 5]. A remodeller (*e.g*., RuvAB, RecG, RecA, RecQ) pushes the fork backwards, allowing the pairing of the nascent strands and rewinding of the parental strands, producing a “chicken-foot” HJ structure [5, 7-10]. In this intermediate, the regressed arm of the reversed fork can be degraded by the RecBCD helicases-nuclease complex (counterpart of *Bacillus subtilis* AddAB), leading to a Y-shaped structure [11, 12]. In addition, the HJ structure can be migrated and cleaved by the RuvABC (counterpart of *B. subtilis* RuvAB-RecU) complex, leading to fork breakage [13, 14]. These dominant mechanisms of fork processing, which use a one-ended double-strand break (DSB) intermediate that will be processed *via* homologous recombination, are instrumental for *E. coli* fork reactivation [15]. Alternatively, the RecQ DNA helicase and the RecJ single strand dependent DNA exonuclease can prevent the accumulation of HJ structures by unwinding and digesting the nascent lagging-strand [16, 17].

The fork breakage mechanism, however, should be lethal during phases where only one copy of the genome is available and homologous recombination cannot occur, as it is the case during the revival of haploid *B. subtilis* spores [reviewed by 18]. These differentiated cells have to exploit other repair sub-pathways to respond in a timely and flexible way to stabilise and remodel a stalled fork, to control fork reversal, to circumvent or bypass the containing-lesion gap by different DNA damage tolerance (DDT) pathways, and indirectly to prevent the formation of deleterious DSBs [4, 19]. Indeed, in the absence of both end resection pathways [*i.e*, in the Δ*recJ* Δ*addAB* strain], which drive the first step of homologous recombination, haploid reviving spores remain recombination proficient and apparently are as capable of repairing ionizing radiation damage as the wild type (*wt*) control [20]. In contrast, spore revival upon DNA damage requires the recombinase RecA, its accessory proteins (*e.g*., RecO, RecR), the DNA translocases (RecG and RuvAB), the DNA helicase RadA/Sms and the DNA damage checkpoint DisA in an otherwise *wt* background [20, 21]. Taking these data into account, the most economical assumption is that these proteins contribute to replication fork remodelling and to the early steps of replication fork rescue, thereby limiting fork breakage. It is known that RecG, RuvAB and RecA enzymes convert a stalled fork into a HJ structure [5, 8, 9]. It would be of significant interest to analyse the role of RadA/Sms and DisA proteins in replication fork remodelling, and their interplay in concert with RecA. (Unless stated otherwise, indicated genes and products are of *B. subtilis* origin).

RecA has been implicated in several steps in response to replication stress, including fork protection, remodelling and restart [3, 8, 20, 22-24]. However, some differences with RecA_*Eco*_ have been reported. First, *in vitro*, RecA, in the ATP bound form (RecA·ATP), nucleates and polymerises on the single-stranded (ss) DNA coated by SsbA only if the RecO positive mediator is present [25], although *in vivo* efficient RecA nucleation also requires RecR [28]. Second, RecA·ATP only catalyses DNA strand exchange in the presence of the two-component (SsbA-RecO) mediator [26, 27]. Finally, a RecA nucleoprotein filament interacts with and loads DisA and RadA/Sms onto a branched intermediate, but DisA or RadA/Sms inhibits the ATPase activity of RecA [29-31]. It is likely that RecA·ATP, in concert with its mediators and modulators, triggers DNA strand exchange. On the other hand, in concert with DisA and/or RadA/Sms, limits the remodelling of stalled replication forks until the lesion is bypassed, and replication re-start until the damage signal is overcome [29]. The latter assumption is consistent with the observation that RecA, in concert with the SsbA-RecO mediator, limits PriA-dependent initiation of DNA replication *in vitro* [20]. Moreover, RecA_*Eco*_ also limits *in vitro* DNA replication [24].

DisA provides a DNA damage checkpoint that delays entry into sporulation and revival of haploid spores until DNA damage has been removed [29, 32, 33]. Here, DisA forms a fast-moving focus that pauses upon exposure to the signal(s) that it recognises [32]. Since DisA pausing requires RecO and RecA, but not AddAB and RecJ [29], the intermediate that DisA recognises should be formed when RecA is engaged with branched intermediates (*e.g*., a stalled fork [an isomer of a displacement loop, D-loop] or a HJ structure). In the absence of DisA, exponentially growing cells remain recombination proficient and apparently are as capable as *wt* cells of repairing DSBs [21, 34, 35]. By contrast, inactivation of *disA* renders exponentially growing cells sensitive to the UV-mimetic 4-nitroquinoline-1-oxide (4NQO), or non-bulky methylating lesions, as the ones generated by methyl methanesulfonate (MMS) [35]. It is likely, therefore, that DisA selectively acts at stalled forks repaired *via* DDT sub-pathways, rather than by canonical DSB repair [21, 35].

DisA, which forms stable octamers, converts a pair of ATPs into a cyclic 3′, 5′-diadenosine monophosphate (c-di-AMP) molecule, an essential second messenger that plays crucial roles in stress management [36]. *In vitro*, DisA bound to stalled or reversed forks reduces c-di-AMP synthesis by 2- to 4-fold [31, 35, 37]. In agreement with this, in response to MMS- or 4NQO-induced lesions, replication stalls, and the amount of the essential c-di-AMP messenger drops by 2-fold in *wt* cells *in vivo*, to levels comparable to that in the absence of DisA [34]. Low c-di-AMP levels increase the production of (p)ppGpp, which in turn inhibits DNA primase and indirectly cell proliferation [38, 39]. It is likely that a fail-safe mechanism to coordinate the cell cycle and maintain cell survival, when there are obstacles that may hinder the progression of the replication fork, is provided by DisA.

Firmicutes RadA/Sms is a hexameric helicase that has been implicated in natural chromosomal transformation and DNA repair. It binds ssDNA and HJ DNA with similar high affinity and unwinds DNA by moving unidirectionally in the 5′ → 3′direction [30, 40]. Moreover, upon interacting with RecA, RadA/Sms also unwinds substrates that cannot process itself [30, 40]. RadA/Sms transiently co-localises in the nucleoid with DisA in ∼30% of unperturbed growing cells [35]. Upon DNA damage, RadA/Sms interacts with DisA and blocks its DAC activity [31, 35].

Previous genetic, cytological and biochemical data support the hypothesis that RecA, DisA and RadA/Sms in concert contribute to prevent replication fork breakage, to protect the stalled or reversed fork, and to circumvent a replicative stress, perhaps *via* different DDT sub-pathways [21, 35]. This is consistent with the following observations: i) DisA or RadA forms a static focus in unperturbed exponentially growing *recU* or *recG* cells, where a significant amount of unresolved branched recombination intermediates accumulate, while they form dynamic foci in *wt* cells [35]; ii) RadA/Sms or DisA limits RecA nucleation and filament growth to prevent RecA from provoking unnecessary recombination during replication fork repair [29-31]; iii) diverse RadA/Sms and DisA mutants directly or indirectly produce dominant-negative phenotypic effects by accumulating toxic DNA intermediates, that are suppressed by *recA* inactivation [29-31]; and iv) DisA or RadA/Sms acts after RecA binding to a lesion-containing gap and prior re-initiation of DNA replication, because DisA neither affects PriA-dependent re-initiation nor DNA replication elongation using a reconstituted *in vitro* DNA replication system [21], but RecA does it [20]. However, the interplay of DisA and RadA/Sms with RecA at the level of the stalled or the reversed replication fork and their contribution to the different DDT sub-pathways remained elusive.

In this work we have investigated this interplay. We show that: i) DisA restrains RadA/Sms and RecA activities; ii) RadA/Sms limits DisA and RecA activities; iii) RecA stimulates DisA and RadA/Sms activities; iv) RecA binds a stalled or reversed fork and facilitates RadA/Sms-mediated reconstitution of the fork to restart replication, but DisA inhibits it; v) RecA reverses the negative effect exerted by RadA/Sms on DisA DAC activity. We propose that fork remodelling is subjected to distinct layers of regulation. RecA at a lesion-containing gap interacts with and loads DisA and RadA/Sms at a stalled or reversed fork, and DisA-mediated c-di-AMP synthesis is suppressed to halt cell proliferation. Then, DisA limits RecA dynamics, and RecA facilitates RadA/Sms unwinding of reversed forks, a reaction limited by DisA. Once the lesion is removed, RecA indirectly antagonises the blockage of cell proliferation by dislodging RadA/Sms, allowing the DAC activity of DisA to be turned on.

## Results

### DisA competes with RadA/Sms C13A for ssDNA

Previously it has been shown that RadA/Sms and its mutant variant RadA/Sms C13A hydrolyse ATP with similar efficiency in the absence of ssDNA [30]. Addition of 3199-nt circular ssDNA (cssDNA) significantly enhanced the rate of ATP hydrolysis of the RadA/Sms C13A mutant variant [29]. RadA/Sms physically interacts with and inhibits the DAC activity of DisA [31]. Both proteins bind stalled or reversed forks and contribute together to maintain genome integrity by a poorly understood mechanism [31]. To understand whether DisA regulates RadA/Sms activities as a part of this crucial interplay, we tested if DisA has any effect on the ATPase activity of RadA/Sms or RadA/Sms C13A.

Accepting that *wt* RadA/Sms or RadA/Sms C13A operates mostly under steady-state conditions, the maximal number of substrate-to-product conversion per unit of time for a 1 μM RadA/Sms or RadA/Sms C13A monomer (k_cat_) was measured. In the absence of cssDNA, the ATPase activity of *wt* RadA/Sms or the RadA/Sms C13A variant (400 nM) (k_cat_ of 9.66 ± 0.2 and 9.60 ± 0.4 min^-1^) was neither stimulated nor impaired by the addition of DisA (500 nM) (k_cat_ of 9.65 ± 0.2 and 9.63 ± 0.2 min^-1^, *p* >0.1) (Fig 1A, green vs yellow line and 1B, orange vs red line). However, the cssDNA-stimulated (10 μM in nt) ATPase activity of RadA/Sms C13A (k_cat_ of 49.1 ± 0.4 min^-1^) was significantly inhibited when DisA was simultaneously added to the reaction (k_cat_ of 20.0 ± 0.5 min^-1^, *p* <0.01) (Fig 1B, yellow vs blue line). Since only the cssDNA-dependent ATPase activity of RadA/Sms C13A is inhibited by DisA, it is likely that DisA might either compete with RadA/Sms C13A for binding to cssDNA, displace it from the ssDNA or inhibit its ssDNA-stimulated ATPase activity by a protein-protein interaction. To evaluate the hypotheses, the effect of the order of protein addition was analysed. When RadA/Sms C13A was pre-incubated with cssDNA (5 min at 37 °C), and then DisA was added, the maximal rate of ATP hydrolysis was moderately reduced (k_cat_ of 32.7 ± 0.2 min^-1^, *p* <0.05) (Fig 1B, brown vs yellow line). On the other hand, if RadA/Sms C13A was added to the preformed DisA-ssDNA complexes, the ATPase activity was further inhibited (k_cat_ of 18.7 ± 0.5 min^-1^, *p*<0.01) (Fig 1B, purple *vs* yellow line). This suggests that DisA competes with RadA/Sms C13A for ssDNA binding.

**Figure 1.**
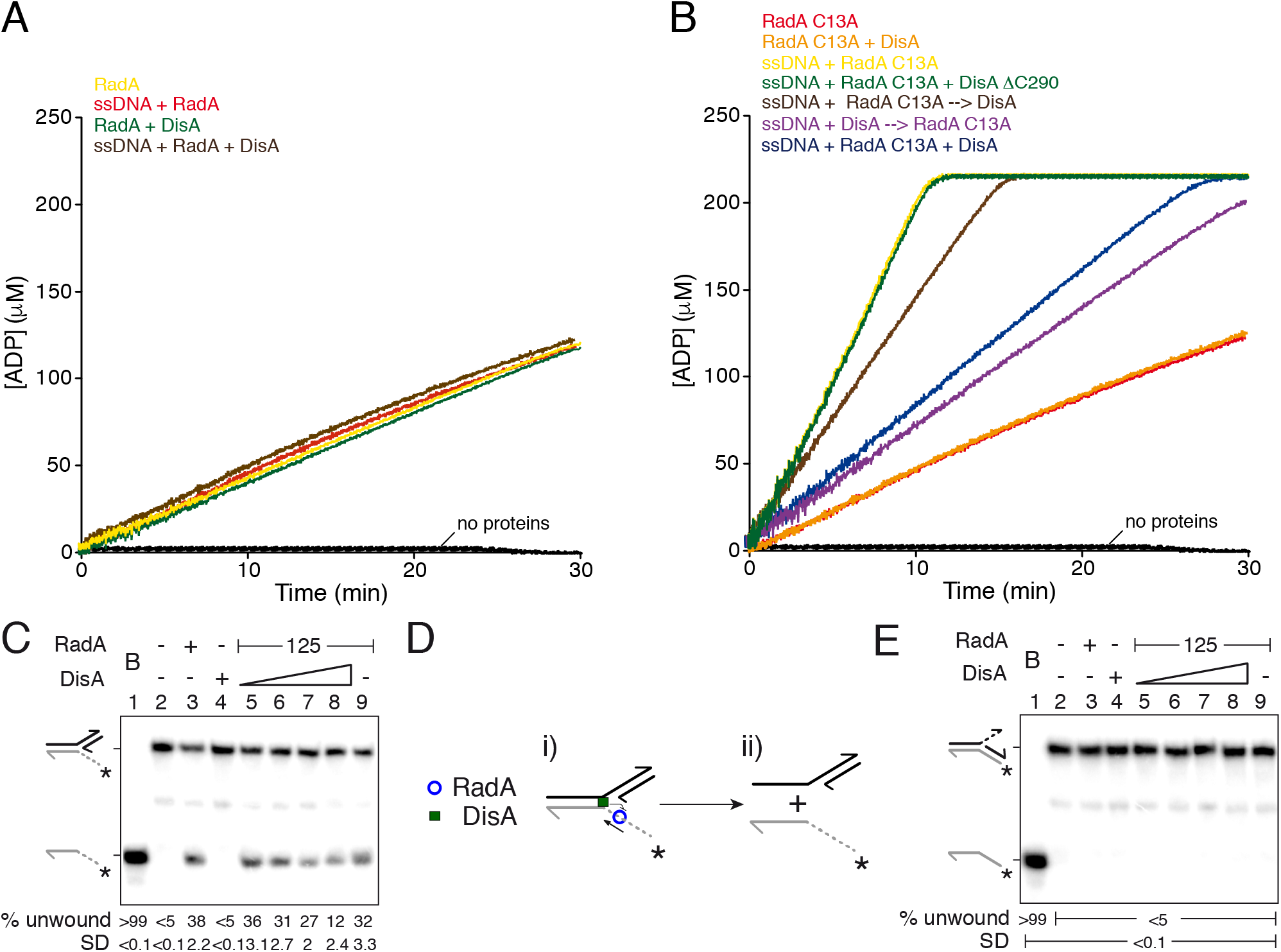
DisA inhibits RadA/Sms activities. (A) RadA/Sms-mediated ATP hydrolysis in the presence of DisA. Reactions had RadA/Sms (400 nM), DisA (500 nM) and when indicated cssDNA (10 μM in nt) in buffer A. (B) cssDNA was incubated with RadA/Sms C13A (400 nM) and DisA or DisA ΔC290 (500 nM), or cssDNA was pre-incubated with RadA/Sms C13A or DisA (5 min at 37 °C), and then DisA or RadA/Sms C13A were added in buffer A. Buffer A contains the ATP regeneration system. (A-B) Reactions were started by addition of ATP (5 mM), and the ATPase activity was measured (30 min at 37 °C). All reactions were repeated three or more times with similar results. A representative graph is shown here, and quantifications of the ATP hydrolysis rates are shown in the main text as the mean ± SD of >3 independent experiments. (C) Helicase assays with 3′-fork DNA. The DNA was incubated with RadA/Sms (125 nM) and increasing concentrations of DisA (100-800 nM). (D) Cartoon illustrating how RadA/Sms unwinds a 3′-fork DNA substrate in the presence of DisA. RadA/Sms unwinds the substrate from its 5′ tail (i-ii), originating a 5′-tailed intermediate, but DisA bound at the junction inhibits RadA/Sms-mediated unwinding. (E) Helicase assays with 5′-fork DNA. The DNA was incubated with RadA/Sms (125 nM) and increasing concentrations of DisA (100-800 nM). (C-E) Reactions were done in buffer A containing 2 mM ATP (15 min, 30°C), and after deproteinization the substrate and products were separated by 6% PAGE and visualised by phosphor imaging. The quantification values of unwound DNA and the SD of >3 independent experiments are documented. Abbreviations: B, boiled DNA substrate; - and +, absence and presence of the indicated protein; * and grey colour, the labelled strand.

To further confirm this conclusion, *wt* DisA was replaced by DisA ΔC290. This mutant variant lacks the DNA binding domain of DisA [37], but still interacts with and its DAC activity is inhibited by RadA/Sms [29]. DisA ΔC290 (500 nM) did not affect the ATP hydrolysis rate of RadA/Sms C13A (k_cat_ of 48.9 ± 0.5 min^-1^, *p* >0.1), when compared to the absence of DisA ΔC290 (Fig 1B, green *vs* yellow line).

In this report, the concentrations of DisA, RadA/Sms and their mutant variants are expressed as moles of monomers, but DisA crystalises as an octamer and RadA/Sms as a hexamer [37, 40]. If we tentatively considered this possibility, it is likely that about equimolar amounts of DisA (1 octamer/∼160-nt, 62 nM) interacts with and competes with RadA/Sms C13A (1 hexamer/∼150-nt, 66 nM) for ssDNA binding.

To test whether DisA bound to cssDNA would inhibit non-specifically the activity of other ATPases at stalled or reversed forks by a competition for DNA binding, the PcrA ATPase was tested. PcrA is another ssDNA-dependent ATPase that acts at stalled or reversed forks and inhibits the ATPase activity of RecA [41]. In the presence of an excess of DisA (800 nM, 1 DisA monomer/ 12-nt), the ATPase activity of PcrA (15 nM, 1 PcrA monomer/ 660-nt) did not significantly vary (k_cat_ of 1750 ± 382 min^-1^ *vs* 1722 ± 332 min^-1^, *p* >0.1) (Fig S1A, red vs blue line). This confirms that the inhibition of the ATPase activity of RadA/Sms C13A by DisA is a genuine and specific activity of DisA.

### DisA cannot activate RadA/Sms to unwind a 5′-fork DNA

Previously it has been shown that Firmicutes RadA/Sms unwinds a 3′-fork DNA (a substrate with a fully synthesised leading-strand and no synthesis in the lagging-strand) by moving in the 5′→3′ direction (Fig 1D) [30, 40]. On the other hand, it cannot unwind a 5′-fork DNA substrate (a fully synthesised lagging-strand and no synthesis in the leading-strand) [30, 40]. However, RecA is sufficient to promote RadA/Sms-mediated unwinding of a 5′-fork DNA substrate, by loading the helicase at appropriate positions on the DNA [30, 40]. Since DisA affects the ssDNA-stimulated ATPase activity of RadA/Sms C13A (Fig 1A-B), to continue studying their interplay, we tested whether DisA regulates RadA/Sms-mediated unwinding using the 3′- or 5′-fork DNA substrates.

Increasing DisA concentrations (100 to 800 nM) significantly reduced (by 2- to 3-fold, *p* <0.05) the unwinding activity of RadA/Sms or RadA/Sms C13A on a 3′-fork DNA substrate (Fig 1C and S2A, lanes 5-8). To test whether this inhibition is solely due to a competition for DNA binding, RadA/Sms-mediated unwinding was analysed in the presence of DisA ΔC290, the mutant variant that cannot bind DNA but still interacts with RadA/Sms [30]. DisA ΔC290 (100-800 nM) inhibited DNA unwinding to a similar extent to *wt* DisA does, with no significant differences observed (*p* >0.1) (Fig S2B, lanes 5-8). This implies that the inhibition of RadA/Sms-mediated unwinding is not caused simply by a competition for DNA binding with DisA, as observed for the ssDNA-dependent ATPase of RadA/Sms C13A. Here, a direct protein-protein interaction or a re-positioning of RadA/Sms on the DNA by DisA might account for this observation.

Finally, we tested whether DisA bound at the junction region of the 5′-fork DNA promotes RadA/Sms loading and RadA/Sms-mediated unwinding of the nascent lagging-strand, as RecA does [30]. Increasing DisA or DisA ΔC290 concentrations did not activate RadA/Sms or RadA/Sms C13A to unwind the 5′-fork DNA substrate (Fig 1E, S2D and S2E, lanes 5-8). This suggests that DisA neither re-positions RadA/Sms to unwind the substrate nor facilitates RadA/Sms-mediated unwinding upon binding at the junction of this substrate.

### RecA-RadA/Sms complexes resist higher ionic strength than RecA-DisA complexes

In previous sections and works it has been defined that a mutual regulation of DisA and RadA/Sms activities exists (Fig 1) [35, 42]. Moreover, it has been postulated that DisA and RadA/Sms separately interact and regulate RecA activities at stalled or reversed forks to maintain genome integrity [29, 30]. In addition, RecA also affects RadA/Sms and DisA activities, as previously introduced [29, 31, 35]. Thus, we investigated the stability of RecA with DisA or RadA/Sms, that is relevant upon DNA damage.

In unperturbed exponentially growing cells, RecA is abundant (∼4000 monomers/colony forming unit [CFU], ∼5.5 μM]), whereas DisA and RadA/Sms are less abundant proteins (∼600 DisA monomers/CFU, ∼800 nM, and ∼500 RadA/Sms monomers/CFU, ∼700 nM) [21, 43, 44]. RecA interacts with both DisA and RadA/Sms proteins [29-31]. Then, before analysing this protein interplay, we have evaluated the strength of such protein-protein interactions.

To perform these experiments, we used native RecA, His-tagged DisA and His-tagged RadA/Sms, and a Ni^2+^ matrix. RecA has a predicted mass of 38.0 kDa, but migrates with an expected mass of 41.5 kDa (Fig S3A, lane 1), and it is not retained in the Ni^2+^ matrix, as described [31]. His-tagged DisA, which has a predicted mass of 40.7 kDa, is contaminated with traces of His-DisA-bound to c-di-AMP (a complex that runs as a 41 kDa protein) (Fig S3A, lane 2), as reported [34].

First, RecA was pre-incubated with His-tagged DisA in the absence of any nucleotide cofactor or DNA (5 min at 37 °C). Then, the mix was loaded onto a 50-μl Ni^2+^ matrix equilibrated with buffer B (Fig S3A). Most of the RecA protein was retained in the Ni^2+^ matrix in the presence of 100 mM NaCl, bound to DisA, and eluted when the matrix was washed with buffer B containing 150 mM NaCl (Fig S3A, lanes 4-5). At 200 mM NaCl, traces of RecA facilitated the release of equimolar amounts of DisA from the matrix (Fig S3A, lane 6). Finally, when DisA bound to the matrix was competitively eluted (E) with buffer B containing 400 mM imidazole and 1 M NaCl, no RecA was observed (Fig S3A, lane 7). In conclusion, it is likely that His-tagged DisA interacts with and retains RecA into the Ni^2+^ matrix, but NaCl concentrations higher than 100 mM are sufficient to disrupt such RecA-DisA interaction.

Similarly, RecA was pre-incubated with His-tagged RadA/Sms (predicted mass 50.3 kDa) in the absence of any nucleotide cofactor or DNA (5 min at 37 °C). Next, the mix was loaded onto a 50-μl Ni^2+^ matrix equilibrated with buffer B. Here, RadA/Sms retained RecA in the Ni^2+^ matrix up to 200 mM NaCl. Last, both RadA/Sms and RecA eluted with buffer B containing 400 mM imidazole and 1 M NaCl (Fig S3B, lanes 4-7). It is likely, therefore, that a higher ionic strength is necessary to disrupt a RadA/Sms-RecA complex, when compared to the DisA-RecA complex.

### DisA and RadA/Sms reduce the ATPase of RecA in a mutually exclusive manner

RecA cooperatively binds ssDNA to form helical nucleoprotein filaments, with a site size of one monomer/ 3-nt [reviewed in 3, 22]. The kinetic of ssDNA-dependent ATP hydrolysis throughout the RecA filament is considered as an indirect readout of its nucleation and polymerization onto cssDNA [3, 22]. RadA/Sms or DisA, which physically interact with RecA (Fig S3), inhibits the ATPase activity of RecA [29, 30]. To begin investigating the global DisA-RadA/Sms-RecA interplay, we tested whether DisA and RadA/Sms in concert regulate RecA nucleation and filament growth using ATPase assays, which provide a real time view of the reaction progress.

Both RecA or RadA/Sms hydrolyses ATP with a k_cat_ of ∼9.6 min^-1^ (Fig 2A, orange and light green lines). The simultaneous addition of cssDNA (10 μM), limiting DisA (100 nM, 1 monomer/100-nt), RadA/Sms (200 nM, 1 monomer/50-nt) and RecA (800 nM, 1 monomer/12.5-nt) significantly blocked the maximum rate of ATP hydrolysis (k_cat_ of 1.0 ± 0.1 min^-1^, *p* <0.01) (Fig 2A yellow line), to levels comparable to the reaction mixture lacking RadA/Sms (Fig 2A, brown *vs* yellow line).

**Figure 2.**
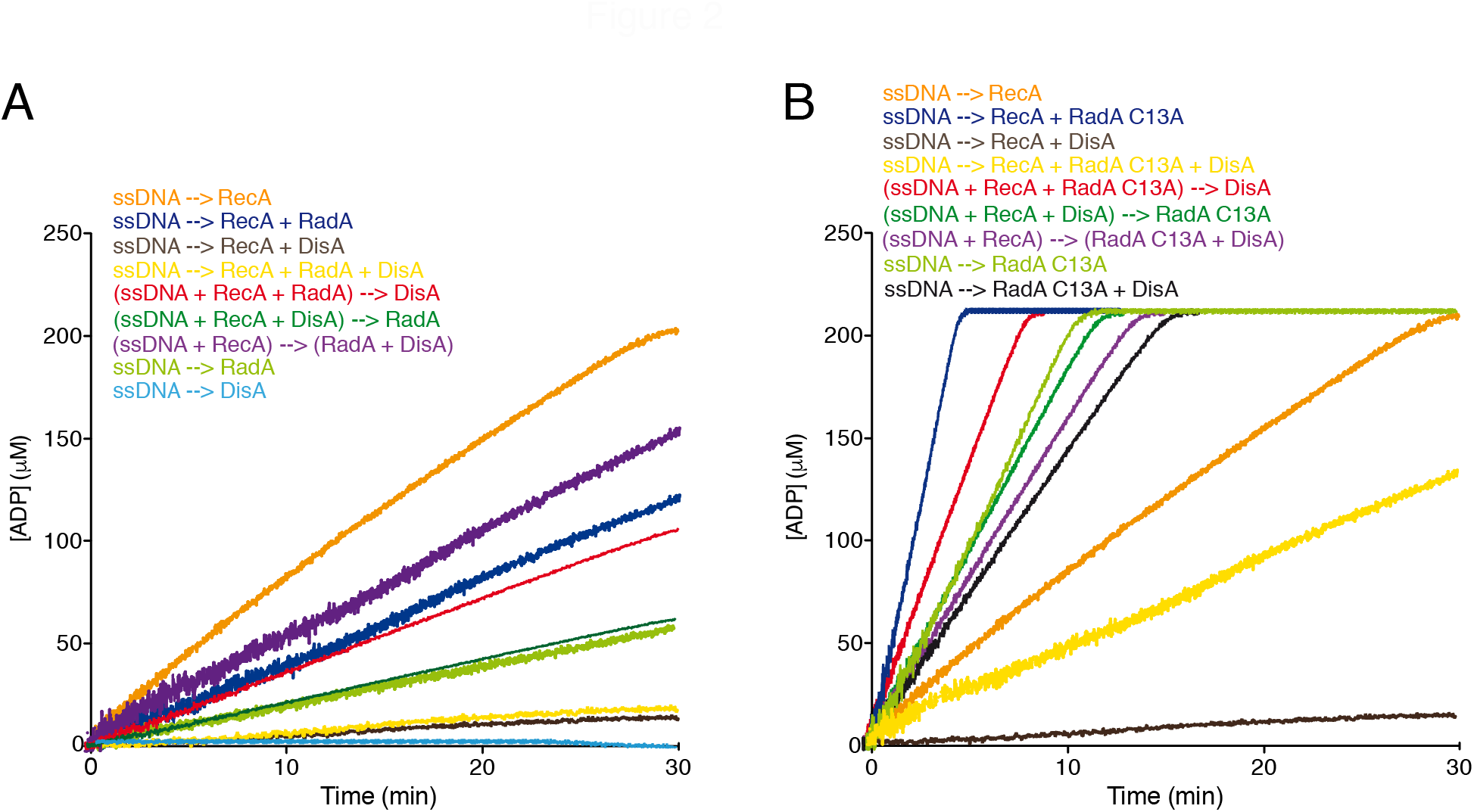
DisA and RadA/Sms competitively reduce RecA-mediated ATP hydrolysis. (A) cssDNA (10 μM, in nt) was incubated with RecA (800 nM), RadA/Sms (200 nM) or DisA (100 nM) or with RecA, RadA/Sms and DisA, or with RecA and RadA/Sms, or with RecA and DisA; or cssDNA was pre-incubated with RecA, or RecA and RadA/Sms or RecA and DisA (5 min at 37°C), then RadA/Sms, DisA or both were added in buffer A. (B) cssDNA was incubated with RecA (800 nM), RadA/Sms C13A (200 nM) or DisA (200 nM), or with RecA, RadA/Sms C13A and DisA, or with RecA and RadA/Sms C13A, or with RecA and DisA, or with RadA/Sms C13A and DisA; or cssDNA was pre-incubated with RecA, or with RecA and RadA/Sms C13A, or with RecA and DisA (5 min at 37°C), and then DisA, RadA/Sms C13A or both were added in buffer A. (A-B) Buffer A contains the ATP regeneration system. Reactions were started by addition of ATP (5 mM), and the ATPase activity was measured (30 min at 37 °C). All reactions were repeated three or more times with similar results. A representative graph is shown here, and quantifications of ATP hydrolysed are shown in the main text as the mean ± SD of >3 independent experiments.

To interpret this regulation, the contribution of the order of protein addition was studied. Addition of DisA to preformed RecA-ssDNA-RadA/Sms complexes reduced the maximal rate of ATP hydrolysis (k_cat_ of 3.5 ± 0.3 min^-1^) to a level comparable to the reaction mixture lacking DisA (k_cat_ of 4.0 ± 0.3 min^-1^) (*p* >0.1) (Fig 2A, red *vs* dark blue line). On the other hand, addition of RadA/Sms to preformed RecA-ssDNA-DisA complexes inhibited the maximal rate of ATP hydrolysis (k_cat_ of 1.7 ± 0.2 min^-1^), but the inhibition was slightly less manifest than when RadA/Sms was omitted (k_cat_ of 0.9 ± 0.1 min^-1^), because RadA/Sms ATPase activity is conserved (Fig 1A *vs* Fig 2A, dark green *vs* brown lines). Finally, when cssDNA was pre-incubated with RecA (5 min at 37 °C), to allow nucleation, and then RadA/Sms and DisA were added, the maximum ATP hydrolysis rate was only moderately reduced (k_cat_ of 5.7 ± 0.3 min^- 1^) (Fig 2A, purple line). It is likely therefore that: i) when the three proteins are incubated together, the activity of both ATPases (RecA and RadA/Sms) becomes inhibited; ii) DisA blocks the ATPase activity of RecA, and addition of RadA/Sms does not reverse this blockage; iii) a preformed RadA/Sms-ssDNA-RecA complex reduces the maximal rate of ATP hydrolysis of RecA, but addition of DisA shows no additive effect; iv) RecA filament growth is less sensitive to the negative effect of RadA/Sms and DisA; and v) the DisA and RadA/Sms activities on RecA-mediated ATP hydrolysis are mutually exclusive.

To further study this protein interplay, RadA/Sms was replaced by the RadA/Sms C13A mutant variant (200 nM, 1 monomer/50-nt), that fails to interact with RecA [30]. In fact, as described [30], when combined with RecA the maximal rate of ATP hydrolysis (k_cat_ of 54.2 ± 0.4 min^-1^, Fig 2B, blue line) approached the sum of the RecA (k_cat_ of 9.6 ± 0.4, orange line) and RadA/Sms C13A (k_cat_ of 49.1 ± 0.4 min^-1^, light green line) independent activities.

When DisA, RadA/Sms C13A and RecA were simultaneously added to cssDNA, the maximum rate of ATP hydrolysis was significantly reduced (k_cat_ of 7.4 ± 0.4 min^-1^, *p* <0.01), but not blocked as it was observed in the absence of RadA/Sms C13A (Fig 2B, yellow *vs* brown lines). This inhibition, however, was ameliorated when limiting DisA was added to preformed RecA-ssDNA-RadA/ Sms C13A complexes (k_cat_ of 30 ± 0.7 min^-1^) (Fig 2B, red line *vs* blue line). Moreover, the inhibition was partially reversed when RadA/Sms C13A was added to preformed DisA-ssDNA-RecA complexes (k_cat_ of 18.2 ± 0.3 min^-1^). Here, the activity was similar to that of RadA/Sms C13A alone (Fig 2B, dark green vs light green lines). Last, when RadA/Sms C13A and DisA were pre-incubated before being added to preformed ssDNA-RecA complexes (k_cat_ of 16.1 ± 0.4 min^-1^), the activity resembles the sum of that of RadA/Sms C13A inhibited by DisA plus that of RecA alone (Fig 2B, purple *vs* black and orange lines). It is likely that DisA interacts with and inhibits the ATPase activity of RadA/Sms C13A and RecA, and both become partially insensitive to DisA action in the presence of the other interacting partner (RecA or RadA/Sms). This confirms the previous observation that DisA and RadA/Sms activities on RecA-mediated ATP hydrolysis are mutually exclusive. Furthermore, the order of protein addition confirms that the DisA inhibitory effect on RadA/Sms and RecA ATPase activity is genuine rather than the presence of a contamination in the protein preparation.

Since DisA synthesises c-di-AMP using ATP as a substrate [35], it may be postulated that the regulation described here is due to a depletion of the ATP pool by DisA or to a direct action of the c-di-AMP molecule over RadA/Sms or RecA. However, we consider this hypothesis unlikely, because it was previously shown that DisA-mediated inhibition is caused by a protein-protein interaction and not by c-di-AMP, using the DisA D77N mutant variant, which does not synthesise c-di-AMP but still inhibits the ATPase activity of RecA [31].

### RecA activates RadA/Sms to unwind 5′-fork DNA, and DisA inhibits it

As previously documented, RadA/Sms is a 5′→3′ DNA helicase [30, 40], but both RecA [30] or DisA (Fig 1C-D) reduces RadA/Sms-mediated unwinding of a 3′-fork DNA substrate. Moreover, RadA/Sms cannot unwind a 5′-fork DNA substrate (Fig 1E), but it can unwind it if RecA is present in the reaction (Fig 3) [30, 40]. To continue understanding the global DisA, RadA/Sms and RecA interplay, we tested whether RecA and DisA in concert regulate the helicase activity of RadA/Sms, using the aforementioned 3′-fork and 5′-fork DNA substrates (Fig 1C and 1E).

**Figure 3.**
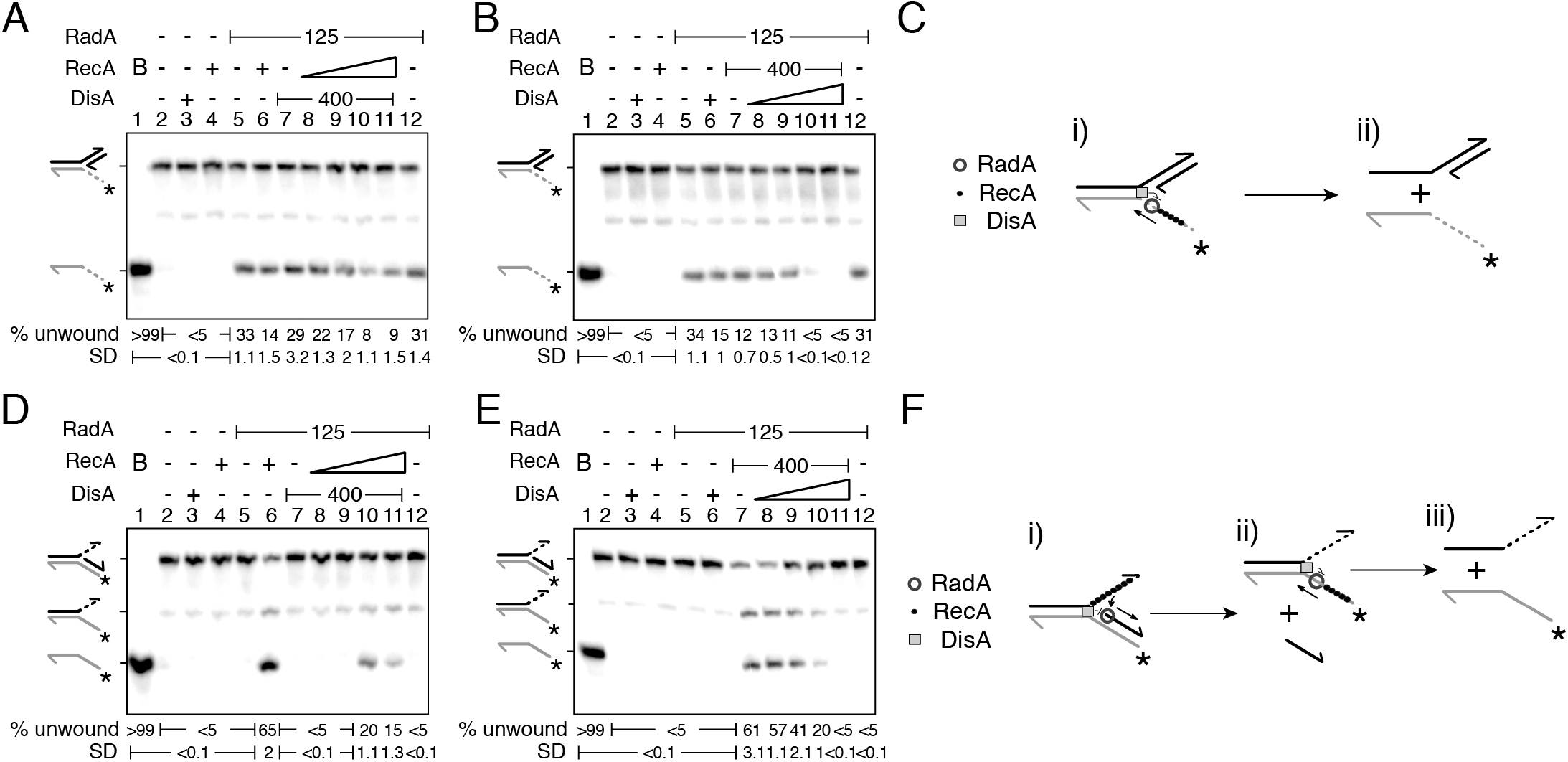
RecA facilitates RadA/Sms unwinding of a 5′-fork DNA substrate, and DisA inhibits it. (A-B) Helicase assays with 3′-fork DNA. 3′-fork DNA was incubated with RadA/Sms (125 nM), increasing concentrations of RecA (50-400 nM) and a fixed amount of DisA (400 nM) (A), or with RecA (400 nM) and increasing concentrations of DisA (100-800 nM) (B). (C) Cartoon illustrating how RadA/Sms unwinds a 3′-fork DNA substrate in the presence of DisA and RecA. RadA/Sms unwinds the substrate from its 5′ tail (i-ii), originating a 5′-tailed intermediate, but DisA bound at the junction and RecA bound at the ssDNA 5′tail inhibit RadA/Sms-mediated unwinding. (D-E) Helicase assays with 5′-fork DNA (). 5′-fork DNA was incubated with RadA/Sms (125 nM), increasing concentrations of RecA (50-400 nM) and a fixed amount of DisA (400 nM) (C), or with RecA (400 nM) and increasing concentrations of DisA (100-800 nM) (D). (C) Cartoon illustrating how RadA/Sms unwinds a 5′-fork DNA substrate in the presence of DisA and RecA. RecA filamented at the ssDNA 3′ tail loads RadA/Sms at the junction on the nascent lagging-strand. Then, RadA/Sms unwinds the substrate (i-ii), originating a forked intermediate and the nascent lagging strand, but DisA bound at the junction inhibit RadA/Sms-mediated unwinding (i-ii). The forked intermediate is further processed by RadA/Sms from its 5′tail, but DisA bound at the junction and RecA bound at the ssDNA 5′tail inhibit RadA/Sms-mediated unwinding (ii-iii). (A-F) Reactions were done in buffer A containing 2 mM ATP (15 min, 30°C), and after deproteinization the substrate and products were separated by 6% PAGE and visualised by phosphor imaging. The quantification values of unwound DNA and the SD of >3 independent experiments are documented. Abbreviations: B, boiled DNA substrate; - and +, absence and presence of the indicated protein; * and grey colour, the labelled strand.

When the 3′-fork DNA was incubated with increasing RecA concentrations (50-400 nM) and fixed amounts of DisA and RadA/Sms, the helicase activity of RadA/Sms was significantly reduced by 4- to 5-fold (*p* <0.01) (Fig 3A, lanes 8-11). However, when the 3′-fork DNA was incubated with fixed RecA and RadA/Sms and increasing DisA (100-800 nM) concentrations, DNA unwinding was blocked at a higher DisA concentration (*p* <0.01) (Fig 3B, lanes 8-11). It is likely therefore that the RadA/Sms:DisA:RecA complex at molar ratios (hexamers:octamers:monomers) of 1:2:20 reduces (by ∼4-fold), and at a molar ratios of 1:4:20 inhibits (by >8-fold) RadA/Sms-mediated unwinding of the 3′-fork DNA substrate. A similar result was obtained when DisA was substituted by DisA ΔC290 (Fig S2C), confirming that DisA-mediated inhibition of RadA/Sms helicase activity is not simply due to competition for DNA binding. Thus, it seems that DisA and RecA affect RadA/Sms-mediated helicase activity in a nearly additive fashion, and the inhibition exerted by each protein can occur at the same time (Fig 3C-D). This agrees with previous results showing that DisA or RecA inhibit RadA/Sms helicase activity *via* different mechanisms: DisA by a protein-protein interaction (Fig 1C and S2), while RecA competes for DNA binding with RadA/Sms, because RadA/Sms C13A-mediated unwinding is also inhibited by RecA, but both proteins do not interact [30].

When the 5′-fork DNA substrate was incubated with a fixed DisA concentration, low RecA concentrations (50 to 100 nM) were not sufficient to activate RadA/Sms-mediated unwinding of a 5′-fork DNA substrate (Fig 3D, lanes 8-9). Indeed, a higher RecA concentration (200 nM) was necessary to activate RadA/Sms unwinding, but in the presence of 400 nM RecA the unwinding reaction was reduced (Fig 3D, lanes 10-11). Under these conditions, RadA/Sms unwound the 5′-fork DNA substrate albeit with ∼4-fold lower efficiency than when DisA was omitted (*p* <0.01) (Fig 3D, lanes 11 *vs* 6).

When the 5′-fork DNA substrate was incubated with fixed RadA/Sms and RecA, and increasing concentrations of DisA (100-800 nM), the RadA/Sms helicase activity was again inhibited in the presence of a molar excess of DisA over RadA/Sms (Fig 3E, lanes 10-11). It is likely that DisA upon interacting with RadA/Sms or with a RecA nucleoprotein filament counteracts the positive effect exerted by the latter on the helicase activity of RadA/Sms. Alternatively, DisA bound at the junction of the 5′-fork DNA competes RecA and abrogates the positive effect exerted by RecA on RadA/Sms helicase activity. To analyse these hypotheses, DisA was replaced by DisA ΔC290. When the 5′-fork DNA was incubated with fixed RadA/Sms and RecA and increasing DisA ΔC290 concentrations (100-800 nM), a molar excess of the latter still counteracted the positive effect produced by RecA over RadA/Sms helicase activity (Fig S2F, lane 10).

Accordingly, it is likely that DisA, by interacting with RadA/Sms or RecA, down regulates RecA-mediated activation of RadA/Sms unwinding of the 5′-fork substrate (Fig 3F). Moreover, an excess of DisA or DisA ΔC290 over RecA inhibits the ability of RadA/Sms to utilise this substrate. It is known that RecA is an abundant protein and lesion-containing gaps increase its expression by 5- to 6-fold as part of the SOS response [44]. Within the *radA* gene maps a stress response promoter for the downstream *disA* gene, whose expression is increased (2- to 3-fold) upon cell envelop stress [45]. Then, we considered that their interplay might in the end limit the activity of RadA/Sms.

### RecA activates RadA/Sms to unwind a reversed fork with a longer nascent leading-strand, but DisA inhibits it

A stalled replicating fork with a leading- or lagging-strand gap may be either spontaneously reversed or remodelled by specialised motor proteins, leading to a HJ-like structure with a longer nascent lagging- or leading-strand, respectively [4, 5]. A HJ-like structure with a longer 5′-nascent lagging-strand can be primed for DNA synthesis and upon fork regression the lesion is bypassed [4, 6]. Under spore revival conditions, however, a HJ-like structure with a longer 3′-nascent leading-strand tail can neither be primed for DNA synthesis nor end resected, because the end resection functions are only synthesised at a later spore outgrowth period [46, 47]. These DNA structures may be processed in two ways. First, re-modellers process the DNA leading to a resetting of the nascent strands back to their original configuration (fork regression) (see Introduction). Second, the 3′-nascent leading-strand tail becomes protected by RecA binding, that it is followed by RadA/Sms-mediated unwinding of the nascent lagging-strand, yielding a 3′-fork DNA (fork restoration) with a longer nascent leading-strand tail (Fig 4Ai-ii).

**Figure 4.**
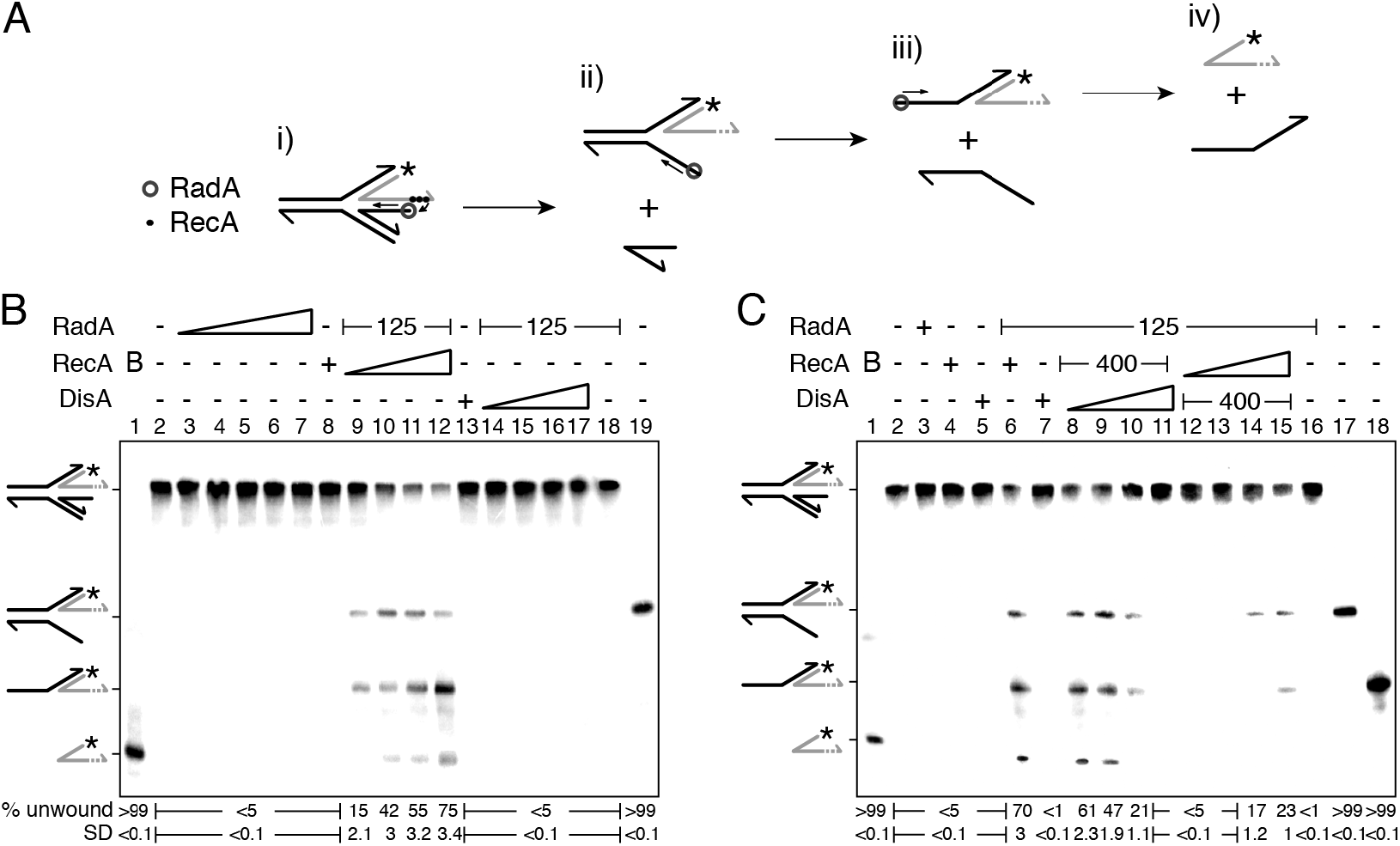
RecA facilitates RadA/Sms-mediated unwinding of a reversed fork with a longer nascent leading-strand, but DisA blocks it. (A) Cartoon illustrating how RecA promotes RadA/Sms-mediated unwinding of 3′-tail HJ DNA substrate. RecA filamented at the nascent leading-strand loads RadA/Sms at the nascent lagging-strand. Then, RadA/Sms unwinds the the newly synthesised lagging-strand (i-ii), originating a 3′-fork intermediate, that is then further processed by RadA/Sms (ii-iv). (B) 3′-tail HJ DNA was incubated with increasing RadA/Sms (30 to 480 nM) concentrations, fixed RecA (400 nM) or DisA (800 nM) concentrations, or with a fixed concentration of RadA (125 nM) and increasing RecA (50-400 nM) or DisA (100-800 nM) concentrations; and the helicase activity measured. (C) 3′-tail HJ DNA was incubated with Rad/Sms (125 nM), RecA (400 nM) or DisA (800 nM), or with a fixed amount of RadA/Sms (125 nM) and RecA (400 nM) and increasing DisA concentrations (100-800 nM), or with a fixed RadA/Sms (125 nM) and DisA (400 nM) and increasing RecA concentrations (50-400 nM); and the helicase activity measured. (B-C) Reactions were done in buffer A containing 2 mM ATP (15 min, 30°C), and after deproteinization the substrate and products were separated by 6% PAGE and visualised by phosphor imaging. The quantification values of unwound DNA and the SD of >3 independent experiments are documented. Abbreviations: B, boiled DNA substrate; - and +, absence and presence of the indicated protein; * and grey colour, the labelled strand.

To investigate this process, an artificial substrate (a HJ-like structure with the nascent leading-strand 30-nt longer than the nascent lagging-strand [3′-tail HJ]) was constructed (Fig 4Ai). This short DNA substrate contains heterologous arms to prevent spontaneous branch migration and the ends of the parental strand are exposed for a secondary action of RadA/Sms (see Fig 4Aiii-iv).

RadA/Sms cannot process blunt-ended HJ structures [30]. In the presence of the 3′-tail HJ DNA and increasing RadA/Sms concentrations (30 to 480 nM), fork regression or fork restoration was not observed (Fig 4B, lanes 3-7). Similar results were observed in the presence of 400 nM RecA or 800 nM DisA (Fig 4B, lanes 8 and 13). In the presence of a fixed RadA/Sms and a limiting RecA (50 nM) concentration, RadA/Sms unwound the nascent and then the parental lagging-strand, yielding a 3′-fork DNA with both a 3′- and a 5′-end available (fork restoration) and a non-replicated fork-like DNA intermediates (Fig 4B, lane 9). Taking into account previous information [30, 40], it is likely that RecA nucleated on the nascent leading-strand of the 3′-tail HJ DNA interacts with and loads RadA/Sms at the junction. Here, RadA/Sms interacts with the 5′-end of the nascent lagging-strand and unwinds it, originating the 3′-fork DNA intermediate (Fig 4Ai-ii). Subsequently, RadA/Sms bound to the 5′-tail of the 3′-fork DNA (Fig 1 and 4B, lane 9), yielding the non-replicated fork-like DNA intermediate (Fig 4Aiii). *In vivo*, the parental strands of a reversed fork have no available ends, but in our short artificial substrate the 5′-end of the 3′-fork is exposed and thus the first intermediate should be unwound by RadA/Sms (Fig 4Aiii-iv). Fork regression or the accumulation of two flapped structures was not observed, suggesting that RadA/Sms cannot regress a 3′-tail HJ DNA.

In the presence of increasing RecA concentrations (100-400 nM), RadA/Sms unwound the 3′-tail HJ DNA, generating the previous two DNA intermediates (Fig 4Aii-iii) and finally the labelled nascent leading-strand (Fig 4Aiii-iv and 4B, lanes 10-12). When RecA was replaced by increasing DisA concentrations (100-800 nM), no unwinding was detected (Fig 4B, lanes 14-18).

To analyse whether DisA affects RecA-mediated activation of RadA/Sms to catalyse branch migration or fork restoration, DisA was added to the reaction (Fig 4C). In the presence of fixed RadA/Sms and RecA concentrations, increasing concentrations of DisA (100-800 nM) significantly inhibited RadA/Sms-mediated unwinding of the 3′-tail HJ (*p* <0.01). Indeed, at a higher DisA molar excess, the unwinding activity of RadA/Sms was finally blocked (Fig 4C, lanes 8-11).

To test whether this blockage is reversible, and if it is due to a competition for ssDNA binding of DisA with RecA, fixed concentrations of RadA/Sms and DisA, and increasing concentrations of RecA (50-400 nM) were used. DisA inhibited RadA/Sms-mediated DNA unwinding of the 3′-tail HJ DNA substrate in the presence of low RecA concentrations (Fig 4C, lanes 12-13). The presence of 200 nM RecA partially counteracted DisA, and RadA/Sms unwound the substrate, yielding the first intermediate (Figs 4Aii) and 4C, lane 14). Last, at 400 nM RecA, the second DNA intermediate was also observed (Fig 4Aiii and 4C, lane 15). It is likely that: i) RecA activates RadA/Sms to unwind the nascent lagging-strand, leading to fork restoration, and DisA downregulates the process; and ii) RecA interacts with and competes DisA.

To test whether the inhibition on the helicase activity of RadA/Sms is a specific activity of DisA, RadA/Sms was replaced by PcrA. In the presence of increasing DisA concentrations (100 to 800 nM), the helicase activity of PcrA (15 nM) did not significantly vary (*p* >0.1) (Fig S1B). This confirms that the inhibition caused by DisA over RadA/Sms-mediated fork restoration is a genuine and specific activity of DisA.

### RecA antagonises RadA/Sms-mediated inhibition of DisA DAC activity

Previously, it has been shown that DisA synthesises c-di-AMP, an essential second messenger. In addition, it has been observed that this synthesis is inhibited when DisA binds DNA branched intermediates, to signal their presence and block cell proliferation until DNA damage is repaired or circumvented (see Introduction). Moreover, RadA/Sms interacts with and inhibits DisA-mediated c-di-AMP synthesis [35, 37]. In previous sections, we have gained insights in the global DisA-RadA/Sms-RecA interplay by analysing the ATPase and helicase activities. Finally, to further understand the protein-protein interactions at stalled or reversed forks, the DAC activity of DisA was measured in the presence of RadA/Sms and RecA.

In the presence of 10 mM Mg^2+^ and 100 μM ATP, DisA converts two ATP molecules into c-di-AMP (K_m_ 151 ± 1.4 μM) [35, 37]. However, this condition is limiting for RadA/Sms-mediated ATP hydrolysis [31]. RadA/Sms or its mutant variants (RadA/Sms K104A or RadA/Sms C13A), above a stoichiometric concentration, interacted with and significantly inhibited DisA-mediated c-di-AMP synthesis (by ∼20-fold, *p* <0.01) (Fig 5A-C, lane 2 *vs* 3).

**Figure 5.**
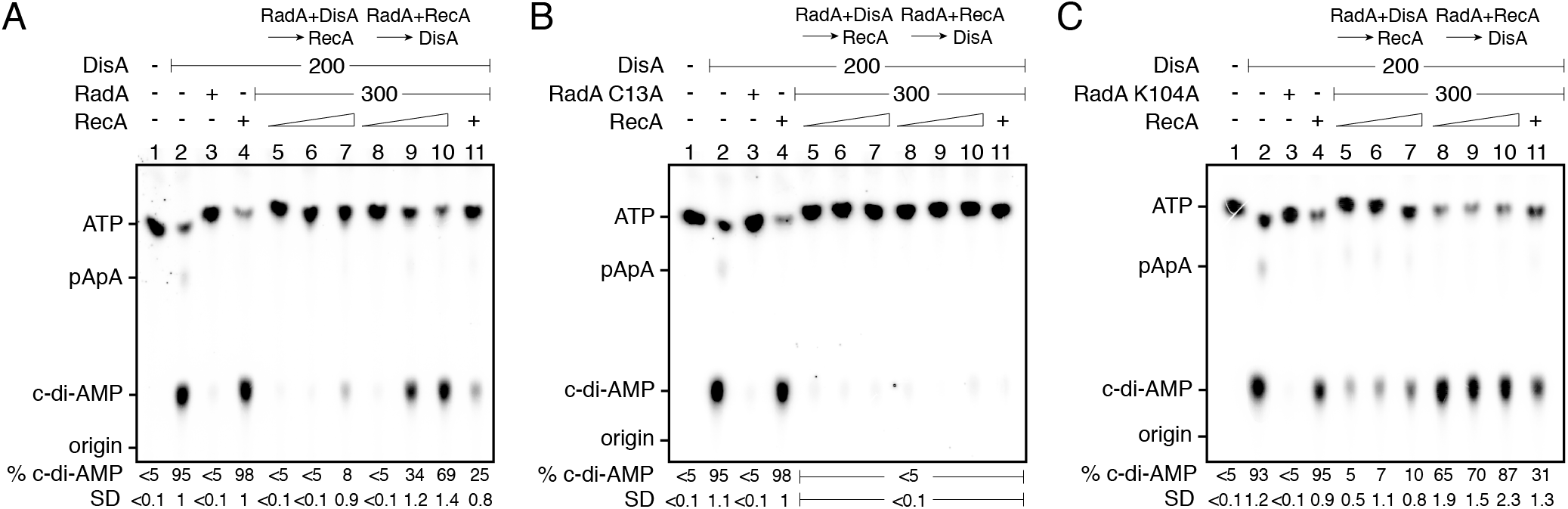
RadA/Sms inhibits DisA DAC activity, but RecA counters this negative effect. (A-C) DisA (200 nM), or DisA (200 nM) and RadA/Sms (A), RadA/Sms C13A (B) or RadA/Sms K104R (C) (300 nM), or DisA (200 nM) and RecA (1600 nM), or DisA (200 nM), RecA (1600 nM) and RadA/Sms (A), RadA/Sms C13A (B), or RadA/Sms K104R (C) (300 nM) were incubated in buffer C containing 100 μM [α^32^P]-ATP:ATP (30 min, 37 °C). DisA (200 nM) was incubated with a fixed concentration of RadA/Sms (A), RadA/Sms C13A (B) or RadA/Sms K104R (C) (300 nM) (5 min, 37 °C), and then increasing RecA concentrations (400-1600 nM) were added in buffer C containing 100 μM [α^32^P]-ATP:ATP (30 min, 37 °C). Fixed RadA/Sms (A), RadA/Sms C13A (B) or RadA/Sms K104R (C) (300 nM) and increasing RecA (400-1600 nM) concentrations were pre-incubated (5 min, 37 °C), and then a fixed amount of DisA (200 nM) was added in buffer C containing 100 μM [α^32^P]-ATP:ATP (30 min, 37 °C). The substrate, intermediates and products were separated by TLC and quantified. The quantification values of c-di-AMP synthesis and the SD of >3 independent experiments are documented. The position of [α^32^P]-ATP:ATP, linear pppA-pA (denoted as pApA), c-di-AMP and the origin are indicated.

An excess of RecA (1600 nM) did not affect DisA-mediated c-di-AMP synthesis (*p* >0.1) (Fig 5A-C, lane 2 *vs* 4). Then, we tested its effect over DisA activity in concert with RadA/Sms. Since the interactions among RecA, RadA/Sms and DisA seem to be mutually exclusive (Fig 2), it can be postulated that RecA bound to RadA/Sms would impede RadA/Sms-DisA interaction and reverse the inhibitory effect of RadA/Sms on c-di-AMP production. To address that, fixed amounts of DisA and RadA/Sms (or its mutant variants) and RecA (1600 nM) were simultaneously added and c-di-AMP synthesis was analysed. Here, RadA/Sms or RadA/Sms K104A only reduced 3- to 4-fold (*p* <0.01) the DAC activity of DisA. However, RadA/Sms C13A blocked the DAC activity of DisA even in the presence of RecA (Fig 5A-C, lane 11), suggesting that a RecA-RadA/Sms interaction is necessary to observe c-di-AMP synthesis restoration. Thus, it is likely that RecA counters RadA/Sms-mediated inhibition of DisA-mediated c-di-AMP synthesis.

To gain insight of the protein-protein interactions, the order of protein addition was analysed. When DisA was pre-incubated with RadA/Sms or RadA/Sms K104A (5 min at 37 °C), and then increasing concentrations of RecA (400-1600 nM) and ATP were added, the DAC activity of DisA was only partially restored at a high RecA concentration (Fig 5A and 5C, lanes 5-7). RecA, however, did not antagonise the preformed RadA/Sms C13A-DisA complexes (Fig 5B, lanes 5-7).

When RadA/Sms or RadA/Sms K104A was pre-incubated with increasing concentrations of RecA (5 min at 37 °C), and then DisA and ATP were added, the preformed RadA/Sms-RecA or RadA/Sms K104A-RecA complex marginally inhibited DisA-mediated c-di-AMP synthesis (*p* <0.01) (Fig 5A and 5C, lanes 8-10). This confirms that RecA antagonises the negative effect of RadA/Sms on DisA-mediated c-di-AMP synthesis. To confirm if the interaction of RecA with RadA/Sms is necessary, RadA/Sms was replaced by the RadA/Sms C13A mutant. When RecA was pre-incubated with RadA/Sms C13A (5 min at 37 °C), and then ATP and DisA and were added, an excess of RecA did not counteract the inhibition of RadA/Sms C13A on DisA-mediated c-di-AMP synthesis (*p* >0.1) (Fig 5B, lanes 8-10).

These data altogether suggest that: i) RecA poorly counteracts the negative effect of RadA/Sms on c-di-AMP synthesis from preformed RadA/Sms-DisA complexes (Fig 5A, lanes 5-7); ii) RecA efficiently counteracts the negative effect of RadA/Sms on DisA-mediated c-di-AMP synthesis when complexed with RadA/Sms (Fig 5A, lanes 8-10); and iii) the RecA-RadA/Sms interaction is necessary to antagonize the negative effect of RadA/Sms on DisA-mediated c-di-AMP synthesis (Fig 5B, lanes 5-11).

### RecA-ssDNA antagonises RadA/Sms-mediated inhibition of DisA DAC activity

Previously, it has been observed that DisA binding to branched intermediates, and in less extent to ssDNA, inhibits c-di-AMP synthesis to signal the presence of DNA damage [35]. RecA efficiently nucleates on ssDNA [3, 22]. Thus, to test whether a preformed RecA nucleoprotein filament also controls the inhibition of c-di-AMP production, the DAC activity of DisA was measured in the presence of ssDNA, RecA and RadA/Sms.

The presence of RadA/Sms, RadA/Sms C13A or ssDNA (10 μM) strongly inhibited DisA-mediated c-di-AMP synthesis (*p* <0.01) (Fig S4A-B, lanes 3-4). On the other hand, a sub-stoichiometric RecA (1600 nM, 1 RecA monomer/6-nt) concentration with respect to ssDNA did not affect DisA-mediated c-di-AMP synthesis (*p* >0.1) (Fig S4A-B, lane 5), but DisA inhibits RecA-mediated ATP hydrolysis (Fig 2) [29]. When DisA was incubated together with RadA/Sms, RecA (1600 nM) and ssDNA, a slight recovery of c-di-AMP synthesis was observed (Fig S4A, lane 12). However, this recovery was not detected when RadA/Sms was replaced by its mutant variant that does not interact with RecA, RadA/Sms C13A (Fig S4B, lane 12).

Last, the effect of the order of protein addition was tested. When increasing concentrations of RecA (400-1600 nM) were added to a preformed RadA/Sms-ssDNA-DisA complex, c-di-AMP synthesis was slightly restored (Fig S4A, lanes 6-8) and to levels similar to that in the absence of ssDNA (Fig 5A, lane 7). Moreover, when RadA/Sms was preincubated with ssDNA and increasing RecA concentrations, and then DisA was added, c-di-AMP synthesis was significantly restored (*p* <0.01) (Fig S4A, lanes 9-11). Nevertheless, when RadA/Sms was replaced by RadA/Sms C13A, RecA failed to antagonize the RadA/Sms C13A inhibitory effect on c-di-AMP synthesis (*p* >0.1) (Fig S4B, lanes 6-11).

It is likely that: i) RadA/Sms, as a part of a preassembled RadA/Sms-ssDNA-RecA complex, cannot exert a negative effect on DisA-mediated c-di-AMP synthesis; and ii) the RecA-RadA/Sms interaction is relevant to the mechanism by which RadA/Sms inhibits the DAC activity of DisA.

## DISCUSSION

Our results support a comprehensive role of DisA in replication fork remodelling and its interplay with RecA and RadA/Sms to prevent their excessive engagement on fork processing, to modulate the choice of DDT sub-pathways and to guarantee population survival. This way, DisA indirectly contributes to the re-establishment of the replication fork and to maintain genome integrity during the revival of haploid spores (Fig 6). Collectively, the study presented here emphasises the importance of timely and flexible responses of DisA to the formation and in the stability of reversed replication forks. Similarly, in eukaryotes, a temporal window is open to allow the access of homologous recombination functions at the stalled fork, and this process is also tightly controlled [6, 19].

**Figure 6.**
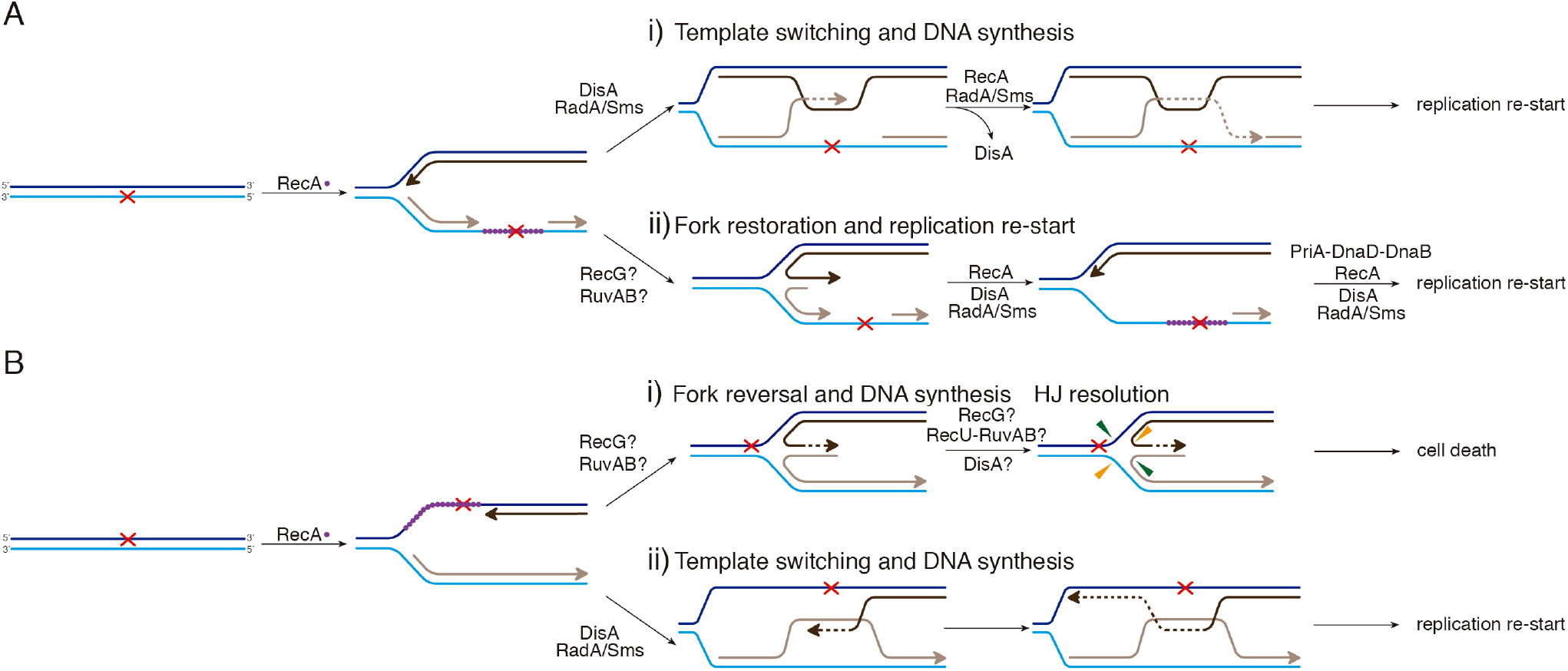
RecA, DisA and RadA/Sms interplay. (A) An unrepaired DNA lesion on the lagging-strand template (red cross) causes blockage of replication fork movement. RecA-bound to the lesion-containing gap suppresses DisA dynamic movement and loads RadA/Sms onto the forked or D-loop structure. (i) RecA promotes D-loop formation and RadA/Sms mediates D-loop extension and the invaded strand primes DNA synthesis. DisA bound to D-loop DNA decreases c-di-AMP synthesis, that in turn increases (p)ppGpp synthesis, that inhibits cell proliferation. DisA suppresses RadA/Sms mediated D-loop extension and RecA reload RadA/Sms in the complementary strand to promote D-loop disassembly. (ii) RadA/Sms unwinds the nascent lagging-strand and allows the fork to be restored. DisA bound to forked DNA decreases c-di-AMP synthesis, that in turn increases (p)ppGpp synthesis, that inhibits cell proliferation. DisA suppresses RadA/Sms mediated DNA unwinding, and DnaD-PriA promote replication re-start. (B) An unrepaired DNA lesion on the leading-strand template (red cross) causes blockage of replication fork movement. RecG or RuvAB branch migration translocases may promote fork reversal. (i) DisA suppresses RecA-mediated reversal of the leading and lagging daughter strands to form a HJ DNA structure. DisA in concert with RuvAB and RecU might contribute to avoid the formation of lethal one-ended DSBs in reviving spores. (ii) RecA-mediated D-loop formation is regulated as depicted in (Ai). DisA bound to D-loop or HJ DNA decreases c-di-AMP synthesis, that in turn increases (p)ppGpp synthesis, that inhibits cell proliferation.

At present, it is unknown the fork remodeller(s) that processes a stalled fork during the revival of *B. subtilis* spores. In *E. coli*, RuvAB, RecG, RecA and RecQ might remodel stalled forks [5, 8, 10]. On the other hand, RecQ and RecJ may prevent the accumulation of HJ structures by unwinding and digesting the nascent lagging-strand [10, 16]. However, when the single genome of an inert mature haploid *B. subtilis* spore is damaged by ionizing radiation, spore revival requires RecA, RecG, RuvAB, RadA/Sms and DisA, but not a RecQ-like DNA helicase (RecS or RecQ) and RecJ functions [20, 21]. Thus, it is likely that RecA, RecG, RuvAB, RadA/Sms and DisA contribute to replication fork remodelling and restart. The role of DisA in limiting RecA activities was previously reported [29], and here, we report the interplay of DisA with RecA and RadA/Sms in the stability of nascent strands after replication fork stalling.

Based on genetic and cytological data, we have inferred that RecA bound to a lesion-containing gap acts prior DisA or RadA/Sms [29, 31, 35]. A RecA nucleoprotein filament interacts with and loads DisA and RadA/Sms at a stalled or reversed fork [29, 30]. It is unknown, however, how these protein-protein interactions support a spatio-temporal regulation, and if they have a role in determining the selection of an appropriate DDT sub-pathway.

On the basis of *in vivo* and *in vitro* data, it is proposed that replicating *B. subtilis* cells often possess only a single active replisome complex, and replisome disassembly and reassembly at each fork is thought to require restarting at least five times per each cell cycle [48]. We propose that when the replisome stalls at or behind the fork, the ssDNA is coated by the essential SsbA protein. The two-component RecO-SsbA mediator contributes to load RecA onto the SsbA-coated lesion-containing gap [27]. Using a reconstituted DNA replication system, it has been shown that RecA in concert with the RecO-SsbA mediator limits PriA-dependent re-initiation of DNA replication [20]. However, in the absence of RecA, the loading of the pre-primosomal protein DnaD is compromised [49]. It is likely that RecA is a two-edged sword and its activities must be regulated.

A correlation between a replicative stress, DisA pausing and RadA/Sms engagement with branched intermediates is suggested. The RecA, DisA and RadA/Sms interactions are mutually exclusive (Figs 1-5), and DisA might limit RecA and RadA/Sms activities (Fig 6A-B). RecA bound to a lesion-containing gap might provoke unnecessary recombination. To avoid RecA-mediated DNA strand exchange occurring at stalled or reversed forks, DisA and RadA/Sms limit the dynamics of RecA, by inhibiting its ATPase activity (Fig 2). If the damage is in the template lagging-strand, RecA invades and pairs the complementary nascent strands, resulting in a D-loop intermediate, in which the nascent leading-strand is used as a template for DNA synthesis of the nascent lagging-strand, contributing to circumvent the DNA damage (template switching) (Fig 6A*i*). However, the D-loop intermediate might be not further extended by RecA. This is consistent with the observation that: i) RecA mediated strand invasion occurs in the absence of ATP hydrolysis, but DNA strand exchange requires ATP hydrolysis for dissociation and subsequent redistribution [3, 22]; and ii) in the absence of the end resection functions (RecQ, RecS, RecJ, AddAB or both RecJ and AddAB), haploid reviving spores remain recombination proficient and apparently are as capable of repairing ionizing radiation damage as the *wt* control [20].

If any of the putative fork remodellers pushes backwards the stalled fork with a lesion in the lagging-strand, it is converted into a HJ-like structure (Fig 6A*ii*) [5, 7, 15]. RecA·ATP filamented on the lesion-containing gap interacts with and loads DisA and RadA/Sms to the reversed fork. Here, DisA has three activities: to suppress RecA dynamics, to limit RadA/Sms-mediated DNA unwinding and to suppress its DAC activity to signal the DNA damage. In the first activity, DisA and RadA/Sms bound to the recombination intermediate interact with and pause RecA dynamics, but the inhibition is not additive (Fig 2). In the second activity, DisA downregulates RadA/Sms-mediated DNA unwinding and RadA/Sms-RecA-mediated fork restoration (Figs 1, 3 and 4). In the third one, DisA bound to a reversed fork, in the presence of RadA/Sms, blocks c-di-AMP synthesis, but this inhibition can be reversed by RecA (Fig 5). Low c-di-AMP levels directly inhibit DNA primase and indirectly cell proliferation [38, 39]. However, DisA neither affects PriA-dependent re-initiation nor DNA replication elongation [21], suggesting that DisA does not act as a protein block to the recruitment of replication proteins and to fork progression.

In addition, we propose that the RadA/Sms-RecA interaction promotes the loading of RadA/Sms at the fork junction. RadA/Sms bound to the nascent lagging-strand catalyses its unwinding to restore a 3′-fork structure (Fig 4), and indirectly prevents fork reversal (Fig 6A*ii*). Then, RecA bound to the 3′-tailed nascent leading-strand and SsbA to the nascent lagging-strand of the restored fork might recruit DnaD and PriA, respectively [49, 50]. The PriA-DnaD-DnaB pre-primosome proteins in concert with DnaI mediate reloading of the replicative DnaC helicase leading to replisome loading and subsequent resumption of DNA replication. Replication re-initiation is directly controlled by RecO-SsbA and RecA, but not by DisA *in vitro* [20, 21, 30]. Finally, RecA, in concert with DisA, competes with and displaces RadA/Sms from the DisA-RadA/Sms-ssDNA (or DisA-RadA/Sms-D-loop) complex (Figs 5 and S4), with free DisA synthesising c-di-AMP to reverse the cell proliferation inhibition [38, 39].

When there is a lesion on the leading-strand template, the stalled fork may be converted into a HJ-like structure with a 5′-tail at the regressed nascent lagging-strand (Fig 6B*i*) [5, 15]. Once fork reversal has occurred and the damage is present in a duplex DNA region, it can be removed by specialised pathways (nucleotide or base excision repair). Here, DisA might block RadA/Sms binding to the longer nascent lagging-strand to cause fork restoration (Fig 1). The nascent leading-strand primes DNA synthesis using the reversed nascent lagging-strand as a template to bypass the lesion after regression. However, the reversed fork might be processed by the RuvAB or RecG branch migration translocase and cleaved by the RecU HJ resolvase, leading to a deleterious fork breakage, that at least in haploid reviving spores induces cell death (Fig 6B*i*). In this pathway, DisA may help to prevent cell death by limiting RecA-mediated fork reversal. The potential contribution of DisA and/or RadA/Sms on RuvAB-RecU or RecG-RecU action will be addressed elsewhere. Alternatively, template switching may occur, with the nascent lagging-strand serving as a template for the nascent leading-strand synthesis (Fig 6B*ii*). Then, upon fork reconstitution, the lesion is on duplex DNA, specialised pathways remove/repair the lesion from the parental strand with DisA protecting genome integrity and synthesising c-di-AMP to indirectly free the DnaG primase [see 39] to re-initiate DNA synthesis if the replisome is loaded.

The mechanism by which fork reversal facilitates replication restart remains unresolved, but it is likely that SsbA and RecA bound to the lesion-containing gap recruit PriA and DnaD, respectively, replication reinitiates, and growth-related processes resume as described above. In *E. coli*, PriA-PriB-DnaT-dependent replication re-start requires restoration of a fork with a nascent leading-strand end in proximity to the junction to facilitate loading of the replicative DnaB (counterpart of *B. subtilis* DnaC) DNA helicase [2-4].

In conclusion, the functional interaction among RecA, DisA and RadA/Sms and their mutual regulation provide bacteria with an essential mechanism that contributes to preserve the nascent DNA at stalled forks. These proteins prevent dangerous cleavage of a reversed fork and DNA breaks, that would be lethal in reviving haploid spores in the absence of an intact sister chromosome. This way, they help to overcome a replicative stress, and promote non-lethal DDT mechanisms such as template switching. The protein interplay described here might apply during exponential growth, since RecA, RadA/Sms and DisA are also necessary here to cope with DNA damage that stalls replication fork progression (see Introduction). It might apply to other bacteria that encode these three proteins too, like *Mycobacterium tuberculosis*, which infects one-third of the world population and cause tuberculosis. Then, understanding the role of RecA, RadA/Sms and DisA in DDT and the enzymes that gain access to a stalled replication fork may provide strong mechanistic basis for DisA or RadA/Sms inhibitors to be used in *Mycobacterium* therapy.

## MATERIALS AND METHODS

### Strains and plasmids

*E. coli* BL21(DE3) [pLysS] cells bearing pCB1020 (*radA*), pCB1037 (*radA*K104A), pCB1035 (*radA*C13A), pCB875 (*disA*), pCB1081 (*disA*ΔC290) and pQE-1 (*pcrA*) genes under the control of a rifampicin-resistant promoter (*P*_T7_) were used to overproduce RadA/Sms (the slash between RadA and Sms names denotes that it has alternative names, the gene is termed *radA*), RadA/Sms K104A, RadA/Sms C13A, DisA, DisA ΔC290 and PcrA proteins, respectively, as described [29-31, 35, 51]. *B. subtilis* BG214 cells bearing the pBT61 (*recA*) plasmid were used to overproduce RecA [52, 53].

### Enzymes, reagents, protein and DNA purification, protein-protein interaction

All chemicals used were analytical grade. IPTG (isopropyl-β-D-thiogalactopyranoside) was from Calbiochem (Darmstadt, Germany), DNA polymerases, DNA restriction enzymes and DNA ligase were from New England Biolabs (Ipswich, MA), and polyethyleneimine, DTT, ATP and dATP were from Sigma (Seelze, Germany). DEAE, Q- and SP-Sepharose were from GE Healthcare (Marlborough, MA), hydroxyapatite was from Bio-Rad (Hercules, CA), phosphocellulose was from Whatman (Maidstone, Kent, UK), and the Ni-column was from Qiagen (Hilden, Germany).

The proteins RadA/Sms (49.4 kDa), RadA/Sms K104A (49.4 kDa), RadA/Sms C13A (49.4 kDa), DisA (40.7 kDa), DisA ΔC290 (33.5 kDa), PcrA (83.5 kDa) and RecA (38.0 kDa) were expressed and purified as described [29-31, 35, 51, 53]. RadA/Sms or DisA and their mutant variants have been purified using the same protocol used for the *wt* protein [29, 31]. Purified DisA shows traces of a slow-moving band of ∼42 kDa that corresponds to c-di-AMP-bound DisA [35]. The purified proteins and its mutant variants lack any protease, exonuclease or endonuclease activity in pGEM3 Zf(+) ssDNA or dsDNA in the presence of 5 mM ATP and 10 mM magnesium acetate (MgOAc). The corresponding molar extinction coefficients for RadA/Sms, DisA, PcrA and RecA were calculated as 24,930; 22,350; 70,375 and 15,200 M^-1^ cm^-1^, respectively, at 280 nm, as described [53]. Protein concentration was determined using the above molar extinction coefficients. The concentrations of DisA (and its mutant variants), RadA/Sms (and its mutant variants), and RecA are expressed as moles of monomers. In this study, experiments were performed under optimal RecA conditions in buffer A (50 mM Tris-HCl pH 7.5, 1 mM DTT, 80 mM NaCl, 10 mM MgOAc, 50 μg/ml bovine serum albumin [BSA] and 5% glycerol).

The nucleotide sequence of the oligonucleotides used is indicated in the 5′→3′polarity: J3-1, CGCAAGCGACAGGAACCTCGAGAAGCTTCCGGTAGCAGCCTGAGCGGTGGTTGAATTCCTCGAGGTTCCTGTCGCTTGCG; J3-2-110, CGCAAGCGACAGGAACCTCGAGGAATTCAACCACCGCTCAACTCAACTGCAGTCTAGACTCGAGGTTCCTGTCGCTTGCGAAGTCTTTCCGGCATCGATCGTAGCTATTT; J3-3, CGCAAGCGACAGGAACCTCGAGTCTAGACTGCAGTTGAGTCCTTGCTAGGACGGATCCCTCGAGGTTCCTGTCGCTTGCG; J3-4, CGCAAGCGACAGGAACCTCGAGGGATCCGTCCTAGCAAGGGGCTGCTACCGGAAGCTTCTCGAGGTTCCTGTCGCTTGCG; 170, AGACGCTGCCGAA TTCTGGCTTGGATCTGATGCTGTCTAGAGGCCTCCACTATGAAATCG; 171, CGATT TCATAGTGGAGGCCTCTAGACAGCA; 173, AGCTCATAGATCGATAGTCTCTAGACAGCATCAGATCCAAGCCAGAATTCGGCAGCGTCT; 172, TGCTGTCTAGAGACTATCGATCTATGAGCT. The 3′-tailed HJ DNA was assembled by annealing J3-1, J3-2-110, J3-3 and J3-4, the 3′-fork DNA by annealing 170, 171 and 173, and the 5′-fork DNA by annealing 170, 172 and 173. The substrates were gel purified as described [54, 55] and stored at 4°C. In the cartoon representation of substrates in Figs 1, 3, 4 and S2, the complementary strands are denoted in solid lines, and the noncomplementary regions in dotted lines. The labelled strand is represented in grey colour. DNA concentrations were established using the molar extinction coefficients of 8780 and 6500 M^-1^ cm^-1^ at 260 nm for ssDNA and dsDNA, respectively, and are expressed as moles of nt.

*In vitro* protein-protein interaction was assayed using His-tagged DisA, His-RadA/Sms and RecA (1.5 μg). Combinations of proteins in buffer B (50 mM Tris-HCl pH 7.5, 50 mM NaCl, 10 mM MgCl_2_, 5% glycerol) containing 20 mM imidazole were loaded onto 50-μl Ni^2+^ microcolumns at room temperature. Then, the Ni^2+^ columns were sequentially washed with buffer B containing increasing concentrations of NaCl (from 100 to 200 mM). Finally, the retained proteins were eluted with 50-μl of Buffer B containing 1 M NaCl and 400 mM imidazole. The proteins were separated by 17.5% (RecA-RadA/Sms) or 10% (RecA-DisA) SDS-PAGE and gels were stained with Coomassie Blue.

### ATP hydrolysis assays

The ATP hydrolysis activity of the RecA or RadA/Sms protein was assayed *via* an NAD/NADH coupled spectrophotometric enzymatic assay [56]. The rate of ATP hydrolysis was measured in buffer A containing 5 mM ATP and an ATP regeneration system (620 μM NADH, 100 U/ml lactic dehydrogenase, 500 U/ml pyruvate kinase, and 2.5 mM phosphoenol-pyruvate) for 30 min at 37°C [56]. The order of addition of circular 3199-nt pGEM3 Zf(+) ssDNA (cssDNA, 10 μM in nucleotides [nt]) and purified proteins is indicated in the text. Data obtained from A_340_ absorbance were converted to ADP concentrations and plotted as a function of time [56]. t-tests were applied to analyse the statistical significance of the data.

### c-di-AMP formation

c-di-AMP formation was analysed using thin-layer chromatography (TLC) and [α-^32^P]-ATP as described [35, 37]. Reactions were performed at 37°C using a range of protein concentrations in buffer C (50 mM Tris-HCl pH 7.5, 50 mM NaCl, 1 mM DTT, 10 mM MgCl_2_, 50 μg/ml BSA, 0.1% Triton, 5% glycerol) containing 100 μM ATP (at a ratio of 1:2000 [α^32^P]-ATP:ATP). The order of addition of circular 3199-nt pGEM3 Zf(+) ssDNA (10 μM in nt) and purified proteins is indicated in the text. After a 30 min incubation, the reactions were stopped by adding 50 mM EDTA. 2 μl of each reaction were spotted onto 20 × 20 cm TLC polyethyleneimine cellulose plates and run for about 2 hours in a TLC chamber containing running buffer D [1:1 (v/v) 1.5 M KH_2_PO_4_ (pH 3.6) and 70% ammonium sulfate]. Dried TLC plates were analysed by phosphor-imaging and spots were quantified using ImageJ (NIH). t-tests were applied to analyse the statistical significance of the data.

### DNA unwinding assays

The different forked DNA substrates used were incubated with increasing concentrations of RadA/Sms or its mutant variants, RecA or DisA, for 15 min at 30°C in buffer A containing 2 mM ATP in a 20-μl volume as previously described [57]. The reactions were deproteinised by phenol-chloroform, DNA substrates and products were precipitated by NaCl and ethanol addition, and subsequently separated using 6% (w/v) PAGE. Gels were run and dried prior to phosphor-imaging analysis, as described above. The bands were quantified using ImageJ (NIH). t-tests were applied to analyse the statistical significance of the data.

**Expanded View** for this article is available online.

## Acknowledgments

The authors thank B. Carrasco and María Moreno-del Alamo for the purification of the RecA and PcrA proteins, respectively, and S. Ayora for comments and proofreading of the manuscript. RT was a PhD fellow of the International Fellowship Program of La Caixa Foundation (La Caixa-CNB). This work was supported in part by Ministerio de Ciencia e Innovación/Agencia Estatal de Investigación (MCI/AEI)/FEDER, EU) PGC2018-097054-B-I00 to J.C.A.

## Author contributions

Conceptualization: RTS and JCA; Investigation: RTS and JCA; Writing-original draft: JCA; Writing-review & editing: RTS and JCA; Supervision: JCA; Funding acquisition: JCA.

