## Supplemental Material for "DisA limits RecA- and RadA/Sms-mediated replication fork remodelling to prevent genome instability"

Figure S1

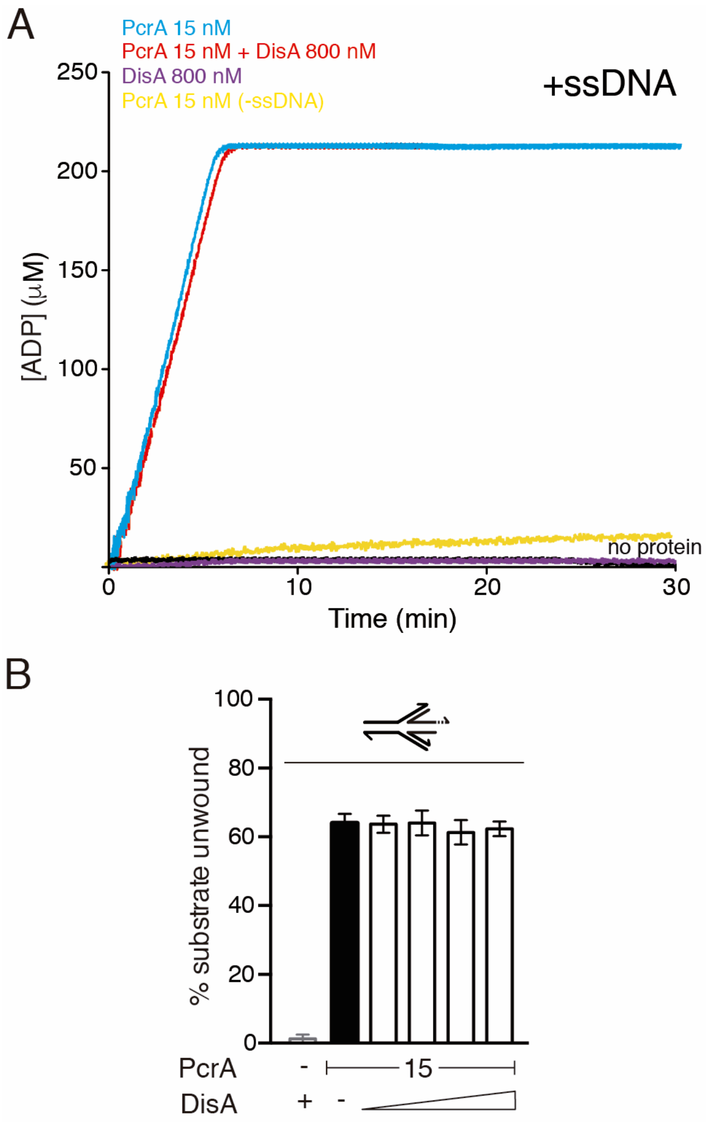


Figure S1. The activities of PcrA are not regulated by DisA. (A) PcrA-mediated ATP hydrolysis in the presence of DisA. Reactions had PcrA (15 nM), DisA (800 nM) and circular 3,199-nt ssDNA (10 μM in nt) in buffer A. Buffer A contains the ATP regeneration system. Reactions were started by addition of ATP (5 mM), and the ATPase activity was measured (30 min at 37 °C). All reactions were repeated three or more times with similar results. A representative graph is shown here, and quantifications of the ATP hydrolysis rates are shown in the main text as the mean ± SD of >3 independent experiments. (B) PcrA-mediated helicase assays with 3´-tail HJ DNA (a HJ-like structure with the nascent leading-strand 30-nt longer than the nascent lagging-strand). The DNA was incubated with PcrA (15 nM) and increasing concentrations of DisA (100-800 nM). Reactions were done in buffer A containing 2 mM ATP (15 min, 30ºC), and after deproteinization the substrate and products were separated by 6% PAGE and visualized by phosphor imaging. The quantification values of unwound DNA and the SD of >3 independent experiments are documented. Abbreviations: B, boiled DNA substrate; - and +, absence and presence of the indicated protein; * and grey colour, the labelled strand.

Figure S2

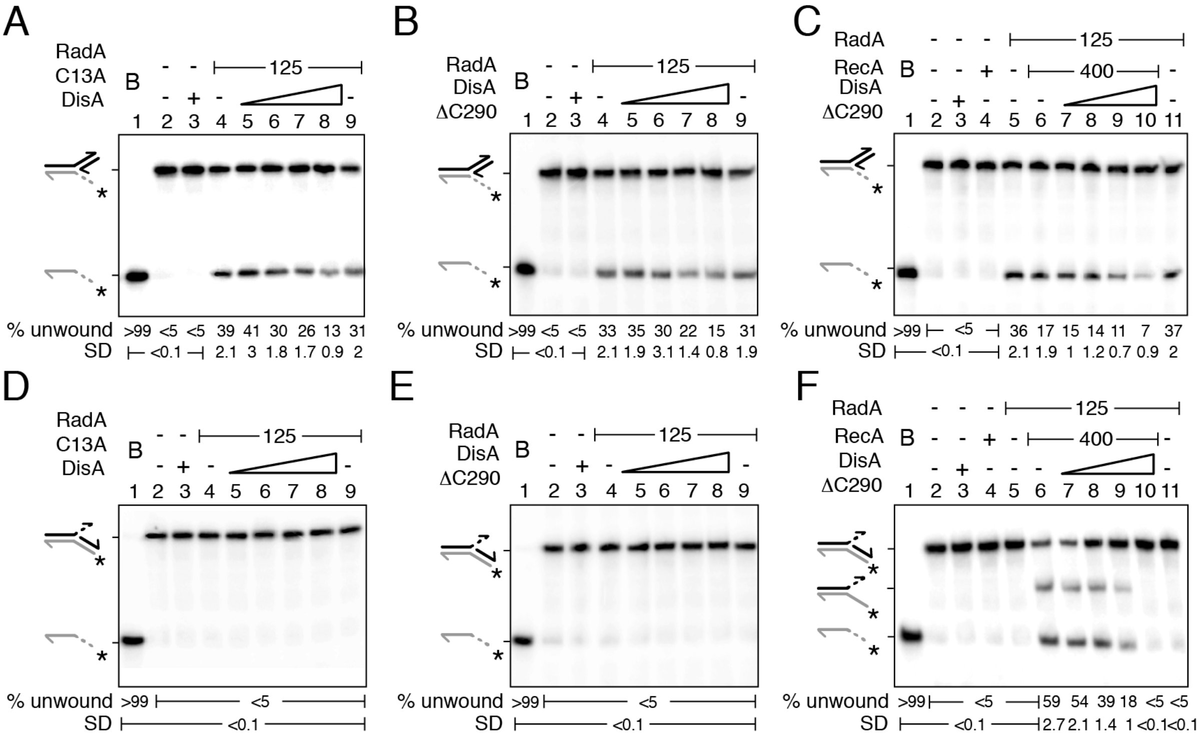


**Figure S2**. DisA action on RadA helicase activity in the presence of RecA. (A-C) DisA ΔC290 partially inhibits RadA/Sms unwinding of 3´-fork DNA. (A) The DNA substrate was incubated with RadA/Sms C13A and increasing concentrations of DisA (100-800 nM). (B) The DNA was incubated with RadA/Sms (125 nM) and increasing concentrations of DisA ΔC290 (100-800 nM). (C) The DNA substrate was incubated with fixed concentrations of RadA/Sms and RecA and increasing concentrations of DisA ΔC290 (100-800 nM). (D-F) RecA loads RadA/Sms to unwind 5´-fork DNA. (D) The DNA was incubated with RadA/Sms C13A (125 nM) and increasing concentrations of DisA (100-800 nM). (E) The DNA substrate was incubated with RadA/Sms and increasing concentrations of DisA ΔC290 (100-800 nM). (F) The DNA substrate was incubated with fixed concentrations of RadA/Sms and RecA and increasing concentrations of DisA ΔC290 (100-800 nM). (A-F) Reactions were done in buffer A containing 2 mM ATP (15 min, 30ºC), and after deproteinization the substrate and products were separated by 6% PAGE and visualized by phosphor imaging. The quantification values of unwound DNA and the SD of >3 independent experiments are documented. Abbreviations: B, boiled DNA substrate; - and +, absence and presence of the indicated protein; * and grey colour, the labelled strand.

Figure S3

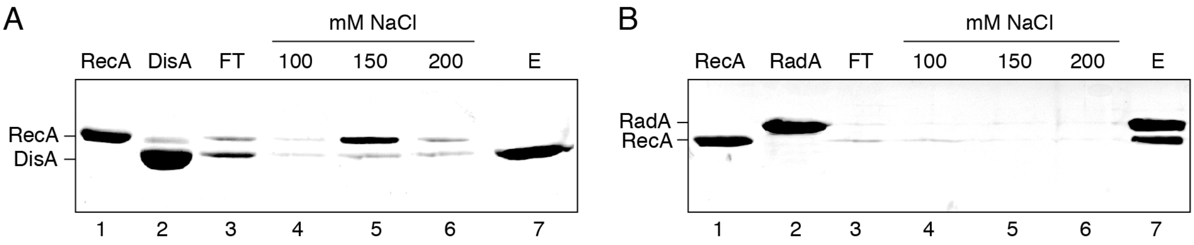


**Figure S3**. RecA forms a stable complex with RadA/Sms, but not with DisA. (A-B) His-tagged DisA (A) or His-tagged RadA/Sms (B) was incubated with RecA in buffer B (50 mM NaCl) (5 min, 37ºC). Then, the mix was loaded onto a 50 µl $Ni^{2+}$ matrix and the flow-through (FT) was collected. The $Ni^{2+}$ matrix was washed once with buffer B and then with 500 µl of buffer B containing increasing NaCl concentrations. Finally, bound His-tagged DisA (A) or His-tagged RadA/Sms (B) was eluted (E) with Buffer B containing 1 M NaCl and 0.4 M imidazole. Experiments were repeated three or more times with similar results, and a representative gel is shown here.

Figure S4

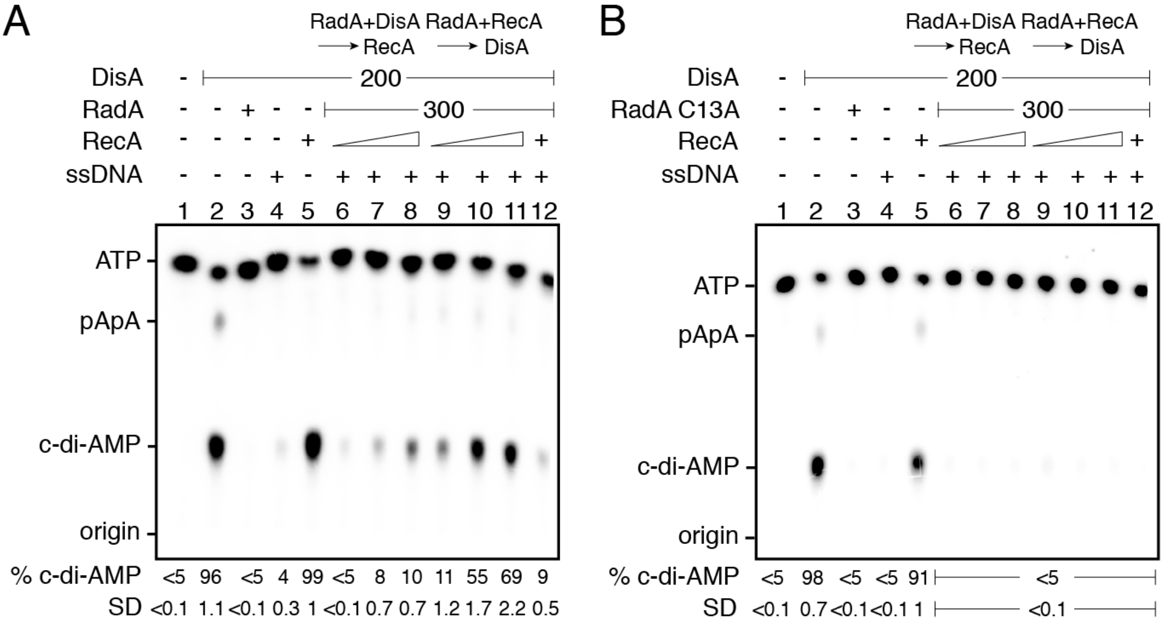


**Figure S4**. RadA/Sms and ssDNA inhibit DisA DAC activity, but RecA counteracts this negative effect. (A-B) DisA (200 nM), or DisA (200 nM) and RadA/Sms (A) or RadA/Sms C13A (B) (300 nM), or DisA (200 nM) and RecA (1600 nM), or DisA (200 nM) and circular ssDNA (10 µM, in nt), or DisA (200 nM), RecA (1600 nM), cssDNA (10 µM, in nt) and RadA/Sms (A) or RadA/Sms C13A were incubated in buffer C containing 100 µM [$\alpha^{32}$P]-ATP:ATP (30 min, 37 ºC). CssDNA and DisA (200 nM) were incubated with a fixed concentration of RadA/Sms (A) or RadA/Sms C13A (B) (300 nM) (5 min, 37 ºC), and then increasing RecA concentrations (400-1600 nM) were added in buffer C containing 100 µM [$\alpha^{32}$P]-ATP:ATP (30 min, 37 ºC). CssDNA, fixed RadA/Sms (A) or RadA/Sms C13A (B) (300 nM), and increasing RecA (400-1600 nM) concentrations were pre-incubated (5 min, 37 ºC), and then a fixed amount of DisA (200 nM) was added in buffer C containing 100 µM [$\alpha^{32}$P]-ATP:ATP (30 min, 37 ºC). The substrate, intermediates and products were separated by TLC and quantified. The quantification values of c-di-AMP synthesis and the SD of >3 independent experiments are documented. The position of [$\alpha^{32}$P]-ATP:ATP, linear pppA-pA (denoted as pApA), c-di-AMP and the origin are indicated.
